# From random to predictive: a context-specific interaction framework improves selection of drug protein-protein interactions for unknown drug pathways

**DOI:** 10.1101/2020.12.15.422844

**Authors:** Jennifer L. Wilson, Alessio Gravina, Kevin Grimes

## Abstract

With high drug attrition, interaction network methods are increasingly attractive as quick and inexpensive methods for prediction of drug safety and efficacy effects when a drug pathway is unknown. However, these methods suffer from high false positive rates for selecting drug phenotypic effects, their performance is often no better than random (AUROC ~0.5), and this limits the use of network methods in regulatory and industrial decision making. In contrast to many network engineering approaches that apply mathematical thresholds to discover phenotype associations, we hypothesized that interaction networks associated with true positive drug phenotypes are context specific. We tested this hypothesis on 16 designated medical event (DMEs) phenotypes which are a subset of adverse events that are of upmost concern to FDA review using a novel data set extracted from drug labels. We demonstrated that context-specific interactions (CSIs) distinguished true from false positive DMEs with an 50% improvement over non-context-specific approaches (AUROC 0.77 compared to 0.51). By reducing false positives, CSI analysis has the potential to advance network techniques to influence decision making in regulatory and industry settings.

**Author summary:** Drugs bind proteins that interact with multiple downstream proteins and these protein networks are responsible for drug efficacy and safety. Protein interaction network methods predict drug effects aggregating information about proteins around drug-binding protein targets. However, many frameworks exist for identifying proteins relevant to a drug’s effect. We consider three frameworks for selecting these proteins and show increased performance from a context-specific approach on selecting proteins relevant to severe drug side effects. The context-specific approach leverages the idea that the proteins responsible for a drug side effect are specific to each side-effect. By discovering the relevant proteins, we can better understand downstream effects of drugs and better anticipate drug side effects for new drugs in development. Further, we focus on designated medical events, a subset of the most severe drug side-effects that are high priority for regulatory review.

## Introduction

Protein interaction network methods are increasingly attractive for understanding drug pharmacodynamic effects because these methods are high-throughput and inexpensive relative to experimental techniques and because they consider drug effects beyond the drug’s targets. Pathways analysis is a valuable tool for understanding a drug’s downstream effects. Yet, curated pathways do not exist for drugs in development and drug pathways do not cleanly align with curated pathways, further necessitating rapid and reliable tools for generating potential pathway mechanisms. To draft these pathways, many have applied network methods for understanding associations between drug target proteins and safety or efficacy phenotypes and extensions of these models have predicted drug repurposing opportunities(1,2) and synergies for drug combinations(3,4). These methods are compelling because a network approach may yield statistically significant associations for a drug’s protein targets to many more phenotypic associations than validated evidence exists. Due to financial or market competition, a drug may only be approved for one, maybe two disease indications, yet a network method might predict statistically significant associations to many more disease indications. These predictions may be opportunities for re-purposing or using drugs off-label yet distinguishing between true positives and true negative disease associations remains a challenge, and gold standard sets of drug effects do not exist(5).

The network community has applied multiple techniques for selecting between true positive and true negative drug phenotypes, each with varying advantages and disadvantages. In an over-simplified view, distance-based methods identify the number of protein-protein interactions to a reach a relevant phenotype association by calibrating to the distance between protein targets of marketed drugs and their intend-to-treat disease genes(1,6,7). Statistical enrichment methods look for enrichment of a network’s genes relative to all gene associations across the entire interactome(2) and some rank drug phenotypes based on the connectivity and closeness of phenotypes shared by drug combinations(8). Neural network methods can achieve high accuracy at labeling known drug-drug interactions using protein-protein interaction networks, drug-target binding data, and gene/protein-phenotype data^4^. However, parsing the potential mechanism behind these predictions remains challenging. Here we compared three different paradigms for separating true positive and true negative drug safety phenotypes to better understand the utility of these paradigms for selecting relevant drug phenotypes.

We specifically considered network associations to drug side-effects because unintended drug side-effects are a major contributor to drug attrition and many applications of network methods have successfully identified drug associations to safety phenotypes. We specifically focused on designated medical events (DMEs) because these are the most severe and are consistently and rigorously considered during regulatory review (e.g. myocardial infarction, pancreatitis) and we did not consider “milder” adverse events (e.g. nausea, rash). By focusing on this subset of adverse events, we reasoned that FDA regulatory review was a sufficiently stringent filter for identifying a true association between a drug and a DME and that a lack of a labeled warning was a sufficient criterion for determining that a drug is *likely* not causative for the DME. Using this assumption, we used a novel data set of positives extracted from drug labels using a natural language processing approach (publication forthcoming). This yielded a set of 1,136 drugs associated to 35 designated medical events (DMEs), a severe subset of drug side-effects. This dataset was originally developed and analyzed to understand patterns in networks of drugs with similar DME associations. However, the dataset provided a unique opportunity to assess the performance of network selection paradigms for identifying relevant drug safety phenotypes using protein-protein interactions. In this dataset, we defined negatives as any of the 1,136 drugs that had network associations to this set of DMEs but the DME was not listed on the drug’s label. We further applied the PathFX algorithm(2) because we could more easily modify the code base to test different paradigms for separating true positives and true negatives.

## Results

### Statistical enrichment cannot clearly separate true positives and true negatives

We first investigated a statistical enrichment method (**Figure 1A**) for separating true positives and true negatives. Specifically, we used PathFX in its original published form. Briefly, PathFX uses a drug’s binding proteins as inputs to identify a network of relevant protein-protein interactions from a larger interactome network (**Supplemental Figure 1**). The algorithm uses a database of gene-phenotype associations and statistical enrichment to identify enriched network phenotypes relative to the original interactome. We used PathFX to identify networks for all 1,136 drugs and investigated where PathFX identified a true positive – a network association between a drug and a DME on the drug’s label – and a false positive – a network association to a DME not listed on the drug label. The distributions for these p-values, both raw and normalized, overlap (**Supplemental Figure 2**), suggesting that a simple statistical test of enrichment is insufficient for separating true positives and true negatives. Not surprisingly, the area under the receiver operator curve (AUROC) is 0.51 (**Figure 1C**).

**Figure 1.**
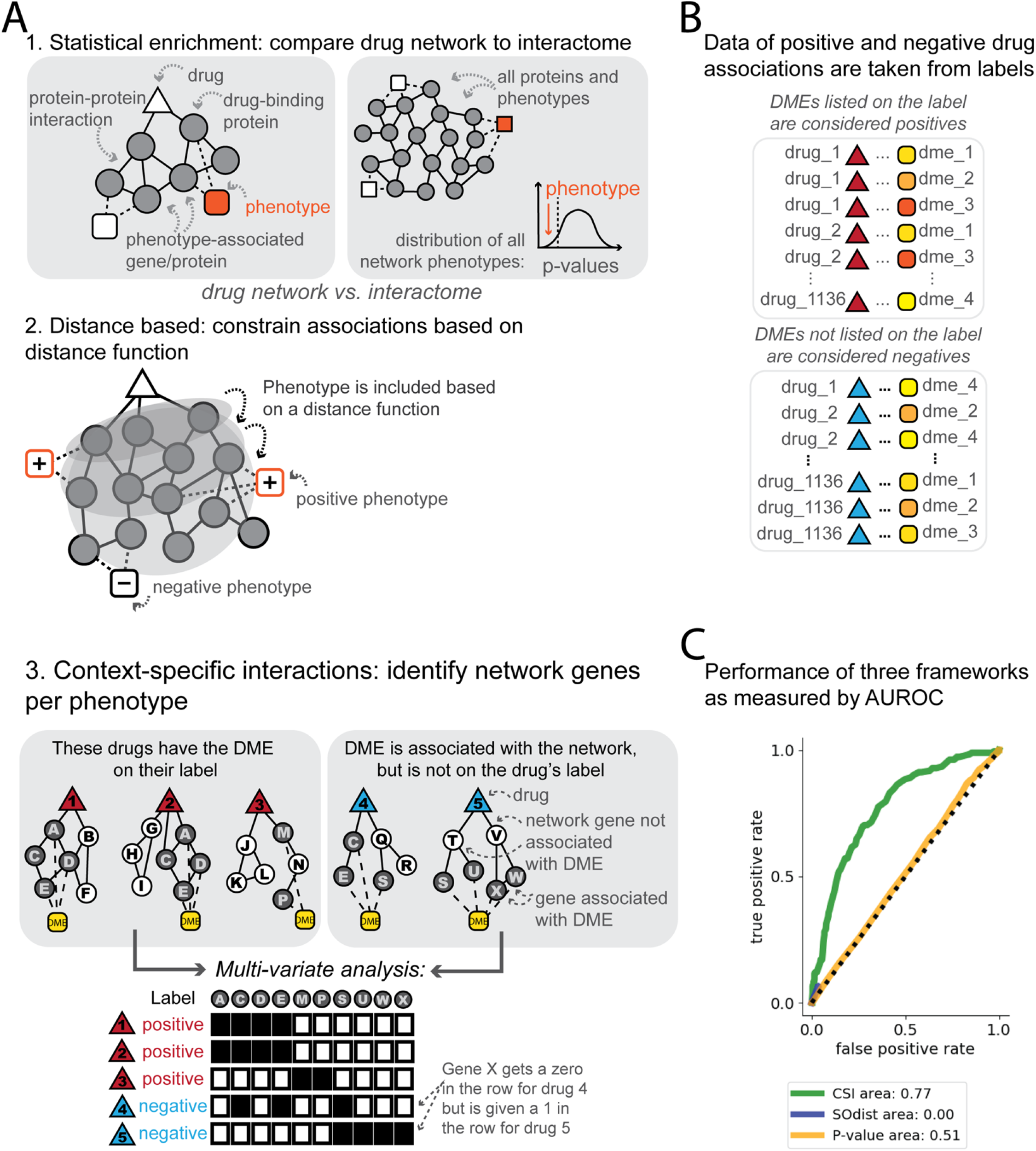
Consideration of three frameworks shows superior performance of context-specific analysis. (**A**) We considered three frameworks: 1. statistical enrichment - a network association is selected if the drug’s interaction network is enriched for associations to a phenotype of interest relative to the entire interactome. 2. distance-based - an interaction distance function is calibrated based on the ability to identify relationships to true positive phenotypes without finding associations to true negative phenotypes. 3. context-specific interactions (CSI) analysis – multivariate analysis (e.g. logistic regression) is used to discover which genes/proteins and interactions separate true from false positives. (**B**) Positive Drug-DME relationships are extracted from the warnings, boxed warnings, and precautions section of the drug’s label. Negative cases (or cases where the drug is not expected to cause the DME) are inferred from the absence of the DME on the drug’s label. Red or blue triangles represent positive or negative drugs, and multiple shades of yellow/orange are meant to distinguish different DMEs in the dataset. (**C**) ROC curves for distinguishing true and false positives using p-value (orange) or a distance-based approach (blue) or using CSIs (green). Legend indicates AUROC value for each framework.

### Using a distance-based approach does not increase model performance for DMEs

We next investigated a simple distance metric for separating true and false positives (**Figure 1A**). For this investigation, we modified PathFX from the original published form (**Supplemental Figure 1)**. Specifically, the original PathFX algorithm relied on an empirically derived path-score threshold to minimize common biases for network algorithms including hub-bias (a gene/protein has high connectivity because it is well studied) and annotation bias (a phenotype is associated with many network genes/proteins because it is overly studied). We considered this path score to be a sufficient proxy for interaction path distance, and so we created modified versions of PathFX using non-optimal distances (e.g. PathFX_dist0.9, PathFX_dist0.8, etc). We reanalyzed our 1,136 drug set using each of these distance algorithms and investigated how relaxing the path score value affected true and false positive rates. At distances of 0.82-0.99, we were unable to generate a full ROC curve(**Figure 1C**). This is likely due to the fact that increasing interaction path distance can only yield more true positives if there are more genes associated with the DME phenotype of interest. We discovered that modifying the path score threshold did not increase an ability to detect true positive associations to DME-associated genes.

### Context-specific interactions increase ability to discern true from false positive DME associations

Much of biology is context dependent and many pathways investigations have used disease-specific pathways to uncover target candidates for therapeutic interventions. We hypothesized that each DME may result from association to a DME-specific pathway and that a better separator of true and false positives could be the specific network genes/proteins supporting an association to a DME phenotype. To test this hypothesis, we tested multiple machine learning and multivariate approaches to distinguish network proteins associated with true positives and true negatives for each DME phenotype. We performed nested cross-validation to select among random forests, logistic regression, and decision trees and used the F1 statistic to discover that these methods were comparable in performance (**Figure 1A, Supplemental Figure 3, Supplemental Table 1**). We selected a simple linear regression because it was the most straightforward method for interpreting if and how network genes/proteins were associated with each DME of interest. Indeed, using a linear regression model combined with networks discovered for DME-associated drugs increased AUROC values 50% improvement over p-value (AUROC 0.77 compared to 0.51) or distance methods (**Figure 1C**). Performance varied for each DME because a separate logistic regression model was required for each DME phenotype (**Supplemental Figure 4**).

CSIs are further attractive for their interpretability. For instance, linear regression feature importance scores highlight network proteins – both drug-binding and downstream of drug-binding proteins – that are associated with positive and negative drugs for each DME (example for edema shown in **Figure 2,** other feature importance scores in **Supplemental File 1**). We overlaid feature importance scores on a merged network for edema to visualize the feature-importance scores in the context of drug protein-protein interaction networks (**Figure 4**). In the tabular results and merged network image, both drug-binding and downstream networked intermediate proteins have high feature importance scores, suggesting that downstream interactions (in addition to specific drug-binding targets) could contribute to drug-induced DMEs.

**Figure 2.**
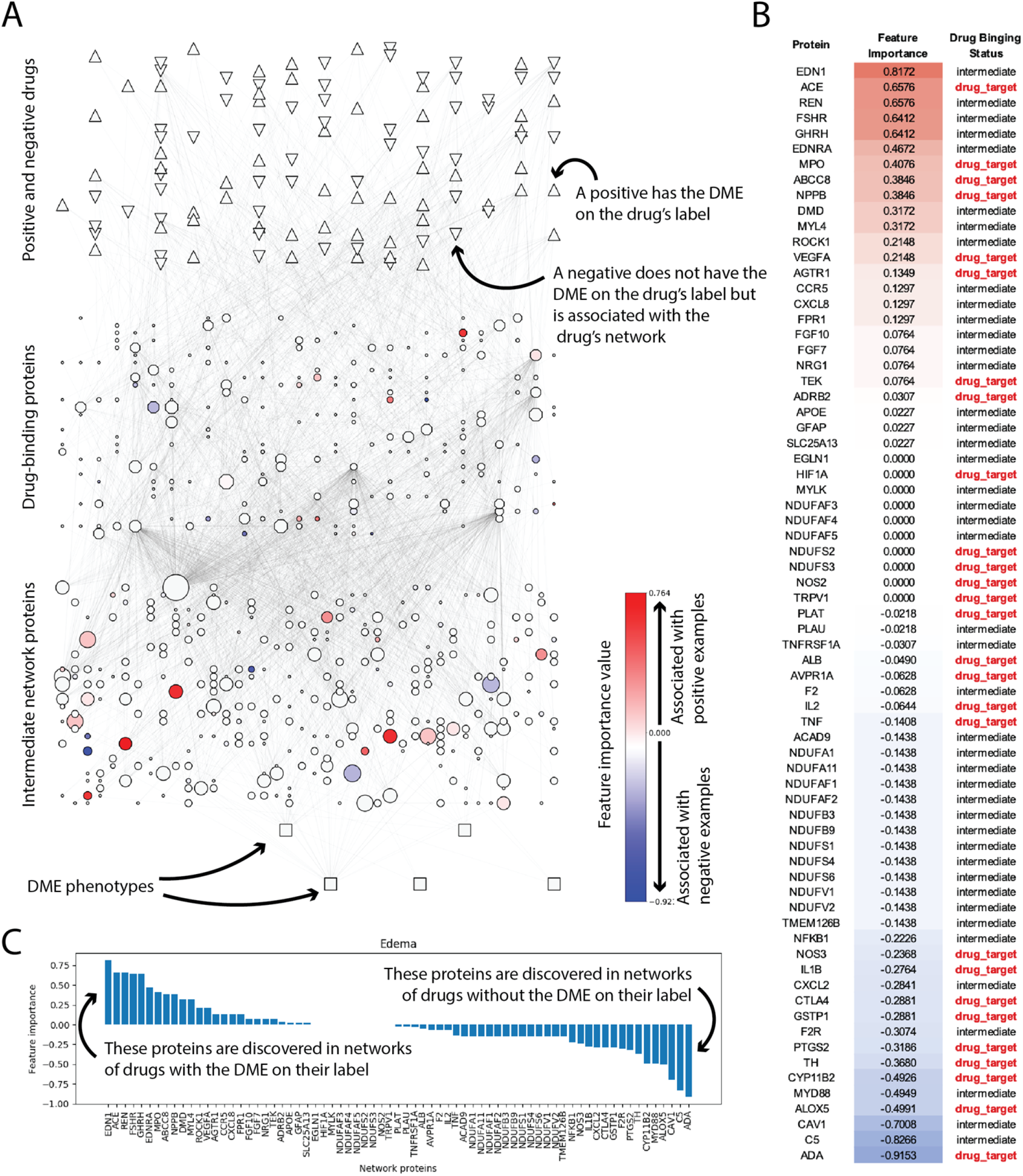
Meta-analysis of DME-associated networks identifies CSIs for edema. The merged interaction network for all true and false positive drugs associated with edema highlights which network components – drug-binding and network proteins – have high feature importance in the logistic regression model (**A**). True/false positive drugs are represented in the top layer as regular/inverted triangles respectively. Drug-binding and intermediate pathway proteins are represented in the second and third layers. The size of the protein reflects the number of networks in which the protein appears. Relevant edema-associated phenotypes are represented as boxes in the last layer. Protein coloring reflects the feature importance in the logistic regression model. Red/blue coloring represents association to true/false positive networks. We have also provided tabular results (**B**) indicating protein feature importance score and whether or not the protein is drug-binding and a histogram (**C**) of ranked feature importance scores.

## Discussion

Protein-protein interaction network methods are increasingly used for identifying phenotypes associated with drug-binding proteins, however, network methods are not sufficiently validated to have translational impact. Here we considered different network selection paradigms for their ability to discern true from false positive drug associations to designated medical events (DMEs). Statistical enrichment is a tractable and relatively easy method to implement, because it requires the selection of a p-value threshold for considering a phenotype as “positive”. However, we discovered that statistical enrichment was unable to separate true positives from true negatives. Distance-based metrics are another attractive, and easily implemented approach for discovering associations between a drug’s targets and DME-associated genes. However, we were unable to universally apply a distance-based metric that correctly identified true positives without increase false positives. Further, interaction distances at high path score thresholds include little to no downstream interactions in the network and these truncated networks can be considered synonymous with only analyzing the drug’s targets. An inability to detect DME associations using only drug targets further motivates the use of network methods for DME detection. We discovered that multivariate and machine learning techniques – specifically a simple logistic regression model – could identify network proteins for each DME and these interaction-based classifiers could separate true positives and true negatives across DMEs. To build further validation and support for network methods to be used more broadly in drug discovery, our results emphasize the importance of leveraging a context-specific paradigm. Indeed, the main contribution of this work is advancing the paradigm of context-specific analysis and emphasizing the role that context-specific interaction “mining” could have for making protein network methods have greater utility in industrial and regulatory decision making.

The relative success of CSI-mining is not entirely surprising given that disease-specific pathway investigations have successfully identified candidate therapeutic targets, however, the results highlight several hypotheses related to advancing network methods to have greater translational impact. In this analysis of DME-associated pathways, it was possible that DME positive and negative drugs converged on the same pathway proteins but had different effects on pathway activation or deactivation. For instance, convergence on the p53 signaling pathway can have both aggravating or mitigating effects on cancer growth depending on the directionality of effect on p53. The superior performance of CSIs suggests that the DME context is important for identifying relevant phenotypes and further, that DME effects could arise, at least partially, from distinct parts of the network – DMEs may arise not from convergence on key network proteins but may arise because of associations to distinct network proteins. Specifically, in the case of edema, our analysis identified endothelin-1 (EDN1) as having a high feature importance score for predicting drugs associated with edema on their labels and EDN1 was not drug binding. In contrast to the p53 example, this highlights a potential downstream signaling effect that could be common to drugs that induce edema. However, further experimental validation would be needed to confirm the relevant of EDN1 or any of the intermediate proteins with high feature importance scores. Across DMEs, we discovered many downstream network proteins, such as EDN1, with high feature importance scores. This further motivates the need to discover and test pathway mechanisms for their role in DME effects in addition to scrutinizing the drug’s direct binding targets. However, the evidence is compelling that context-specific interactions are a complementary and viable paradigm for advancing network methods to have greater influence on decision making in industry and regulatory settings.

We acknowledge limitations in this analysis. For instance, the definition of a set of gold-standard true negative examples is imperfect. We considered the lack of a warning on a drug’s label as a sufficient standard for our analysis, considering the rigor and integrity of the FDA review process. Yet, it’s still possible that some of our true negatives are false negatives and the negative drugs could have a meaningful association to a DME despite a lack of a labeled warning. Because defining gold standard true negatives is difficult, our analysis is limited to the investigation of DMEs and does not consider drug efficacy or milder drug side effects. Understanding how well interaction pathways associate drug targets to efficacy phenotypes would require greater transparency about ‘failed’ tests of drugs against multiple diseases and better curation of this type of data. Further, the current definitions of CSIs could be further optimized. For some DME contexts, we achieved AUROC values that were much higher than p-value or distance-based metrics. For some DME contexts, we were unable to build sufficiently predictive models and ultimately restricted analysis to DMEs where we had at least 10 positive and negative examples. These models could be improved by further curation of positive and negative examples or the inclusion of more data (e.g. drug-protein binding data, gene-DME associations, or protein-protein interactions).

Further advancement of context-specific network analysis will require a sufficient number of known effectors. Specifically, to associate a drug with a relevant effect on a DME, we needed several examples of drugs that cause the DME and drugs that had network associations to the DME but did not cause it. Because we lack these positive and negative examples for understudied diseases without current therapies, we are unable to sufficiently predict drug effects on these understudied diseases. In the case of DMEs, sufficient drug effectors existed for training classifiers, making context-specific analysis feasible for classifying the effects of new compounds on DMEs. We anticipate that data from established assays and model systems will be essential to mining CSIs for drug-efficacy related phenotypes, especially for disease areas where successful therapies do not yet exist. We anticipate that network selection methods relying on mathematical principles (e.g. p-value selection, network distance, network connectivity) will remain as powerful workhorse techniques, especially in contexts where known drug effectors are not established. We see CSI mining as a means to advance these predictions towards the goal of providing predictive power for decision-making at the level or regulatory review or industrial selection of gene targets for new therapies.

The contribution in this work is the demonstration that prioritizing specific subsets of the interaction network can be predictive for modeling drug effects. From a network engineering perspective, CSI mining may indeed affect how network methods are developed for understanding drug mechanisms. The result presented here encouraged us to pursue further validation of the prioritized network proteins as mediators of drug mechanisms. Indeed, combination drugs that bind DME-network proteins synergized to affect adverse outcomes associated with drugs associated with the DME on their drug labels (in preparation). Together, these results suggest a paradigm shift towards network engineering of context-specific pathways to identify drug network mechanisms.

## Materials and Methods

### Extracting true positive and true negative drug examples from drug labels

We extracted relevant phenotypes from the drug’s labels using a custom NLP query (publication forthcoming, data included in *Drugs_labeled_for_AEs.txt*). We further refer to ‘positive drug examples’ as those drugs associated with a DME on their drug label. We refer to ‘negative drug examples’ as any of the 1,136 drugs in our drug set that do not have a DME listed on their drug label. We define positives and negatives for each DME separately. For instance, 496 drugs are associated with myocardial infarction on their drug label, and these drugs are considered positives for the myocardial infarction DMEs. The remaining 640 drugs in our 1,136 drug set are considered negatives for the myocardial infarction DME.

### Modeling true positive and negative networks with PathFX

We analyzed 1,136 drugs using PathFX with default parameter settings. Briefly, PathFX uses drug-binding proteins as inputs to first identify a relevant protein-protein interaction network around these targets, and second uses the full list of network genes/proteins to identify phenotypes associated with these genes/proteins relative to the entire interactome. For each drug, PathFX analysis yielded interaction networks and a list of phenotypes associated with these networks. For a full list of features and outputs, see^2,9^.

We assessed whether a phenotype matched a DME from the drug label and considered these pairs as true positives. We also searched these same association tables for DMEs not listed on the drug’s label and considered these as false positives. The script *define_tp_fp.py* creates the following outputs: *drugs_to_dmes_true_positive.txt, drugs_to_dmes_false_positives.txt*.

### Plotting p-value distributions and estimating AUROC values

In the same script (*define_tp_fp.py*) where we defined our true and false positive examples, we generated plot of the p-values for these associations. This script generates the plots, *raw_pvalues.png*, and *norm_pvalues.png*, and generates the data object, *pvalue_roc_values.pkl*, for further analysis. We analyzed the AUROC in the script, *plot_ROC_pv_soDist.py*, using the trapez method implemented in Python.

### Measuring the effect of interactome distance on detecting DME associations

We developed modified versions of PathFX to test the effect of altering interaction distance on detecting associations to DMEs. For the original PathFX construction we empirically derived an interaction score threshold to prevent hub bias(2). To measure the effect of interactome distance on detecting associations, we created 11 custom versions of PathFX. These scripts are contained in the *PathFX_soDist/scripts/* directory and are named *phenotype_enrichment_pathway_so_dist_0.82.py* where ‘0.82’ represents the score threshold used in this version. The other score thresholds used include 0.82-0.90, 0.95, and 0.99. We used these thresholds out of convenience. We started our experiment using a stringent, high threshold, and then relaxed this threshold to increasingly allow more edges to be considered in network construction. Given the score distribution of our interaction network, we found that computational time increased significantly as we reached the 0.82 range. We truncated the analysis with this version because we saw that we were no longer drastically changing our ability to detect more true positives.

We created networks for our 1,136 drug set using each version using the script *run_pathfx_all_distances.py*. This script generated networks, and association files for all 1,136 drugs at each of the 11 distances. We investigated whether the networks for these drugs contained associations to true or false positive DMEs at each score threshold. We analyzed these results using the script, *count_tp_fp_so_dist.py*, and the results of this script are saved in the *PathFX_soDist/results/analyze_so_dists/* directory. We then used the *plot_ROC_pv_soDist.py* to count the true and false positive rates at each score threshold and plot the ROC curve using these values. This script generates *the pvalue_ROC_dist.png* figure.

### Logistic regression, decision trees, and random forests analysis

For this analysis, we created binary matrices for all true and false positive networks associated with a DME. These matrices included a 1/0 if a gene was/was not included in the drug’s network respectively. Rows were labels as positive if the drug’s label included the DME on the label or negative if the drug’s label was not associated with the DME. This analysis is included in the script *create_positive_negative_files.ipynb* and this analysis yielded a matrix file for each of 24 DMEs: agranulocytosis, cardiac arrest, cerebral infarction, deep vein thrombosis, delirium, edema, gastric ulcer, hemolytic anemia, hemorrhage, hepatic necrosis, hyperlipidemia, hypertension, interstitial lung disease, myocardial infarction, myopathy, pancreatitis, peripheral neuropathy, pneumonia, proteinuria, pulmonary edema, sepsis, tardive dyskinesia, thrombocytopenia, and ventricular tachycardia. These files are saved in */ML_network_positives_negatives/dme_DMENAME.txt* where DMENAME is replaced with each of the DMES of interest.

We first used the scikit-learn module in python and a nested cross-validation procedure to evaluate modeling types – logistic regression, decision trees, and random forest – and used the F1 score to evaluate model performance in this analysis (*ML_network_positives_negatives/run_all_dme_models_ncv.py*). The results of those analysis are included in **Supplementary Figure 1**. To generate test scores for the ROC curves using logistic regression, we modified *ML_network_positives_negatives /run_all_dme_models_new_log_reg.py* and *ML_network_positives_negatives/all_pathways.py* scripts. To plot all ROC curves, we used *plot_ROC_all_methods_072720.py*.

### Plotting merged networks

To analyze feature importance scores, we used *ML_network_positives_negatives/save_and_plot_feat_imp_scores.py*. This script analyzed the feature importance scores generated after the model fitting and generated **Supplemental File 1.** This file is a copy of *ML_network_positives_negatives/log_reg/logistic_regression_all_feature_importance.xlsx*. We next plotted merged networks and feature importance values using *network_images/ plot_feat_imp_on_networks.ipynb*.

## Data and code availability

The data and code used in this analysis are available on GitHub (https://github.com/jenwilson521/network_selection).

## Acknowledgements

JLW was partially and AG and KG were supported by the SPARK program at Stanford. JLW was also supported by a Sanofi iDEA Award.

## Author Contributions

JLW conceived of the idea. JLW and AG conducted the analysis. JLW wrote the manuscript. JLW, AG, and KG revised the manuscript.

